# Improvising at Rest: Differentiating Jazz and Classical Music Training with Resting State Functional Connectivity

**DOI:** 10.1101/732073

**Authors:** Alexander Belden, Tima Zeng, Emily Przysinda, Sheeba Arnold Anteraper, Susan Whitfield-Gabrieli, Psyche Loui

## Abstract

Jazz improvisation offers a model for creative cognition, as it involves the real-time creation of a novel, information-rich product. Previous research has shown that when musicians improvise, they recruit regions in the Default Mode Network (DMN) and Executive Control Network (ECN). Here, we ask whether these findings from task-fMRI studies might extend to intrinsic differences in resting state functional connectivity. We compared Improvising musicians, Classical musicians, and Minimally Musically Trained (MMT) controls in seed-based functional connectivity and network analyses in resting state functional MRI. We also examined the functional correlates of behavioral performance in musical improvisation and divergent thinking. Seed-based analysis consistently showed higher connectivity in ventral DMN (vDMN) and bilateral ECN in both groups of musically trained individuals as compared to MMT controls, with additional group differences in primary visual network, precuneus network, and posterior salience network. In particular, primary visual network connectivity to DMN and ECN was highest in Improvisational musicians, whereas within-network connectivity of vDMN and precuneus network was higher in both Improvisational and Classical musicians than in MMT controls; in contrast, connectivity between posterior salience network and superior parietal lobule was highest in Classical musicians. Furthermore, graph-theoretical analysis indicated heightened betweenness centrality, clustering, and local efficiency in Classical musicians. Taken together, results suggest that heightened functional connectivity among musicians can be explained by higher within-network connectivity (more tight-knit cortical networks) in Classical musicians, as opposed to more disperse, globally-connected cortical networks in Improvisational musicians.

**Highlights:**

- Music training is associated with higher resting state connectivity
- Higher connectivity in Improvisational musicians from visual network to ECN and DMN
- Classical musicians show higher vDMN and Precuneus within-network connectivity
- Improvisation and divergent thinking performance correlate with similar connectivity patterns

## 1. Introduction

Creativity is commonly defined as “the ability to produce work that is both novel (i.e. original, unexpected) and appropriate (i.e. useful, adaptive concerning task constraints)" (Sternberg, 1999, p. 3). Creativity researchers have long wondered the following questions: 1) How do people create novel and appropriate works? 2) What neural systems are recruited for such processing? and 3) Why do individuals differ in creative ability? In addressing the first of these questions within the study of musical improvisation, which is a real-time creative behavior (Loui, 2018), behavioral models suggest that musical improvisation requires a balance between automatic retrieval of candidate musical sequences from long term memory, and continuous appraisal of these sequences over the course of an improvisational performance (Pressing, 1988). This interplay of generative and evaluative behaviors is also posited in other domains of creativity, such as visual art (Ellamil et al., 2012) and creative writing (Liu et al., 2015), and is the basis of dual-process theories of cognition towards creative behavior (Sowden, Pringle, & Gabora, 2015).

Recent work has identified the default mode network (DMN) and the executive control network (ECN, similar to the frontoparietal network FPN), as cortical hubs that may underlie the dual processes of creative cognition (Beaty et al., 2015; Beaty et al., 2018). The DMN, which has been closely associated with self-referential processing and mental simulations (Kim & Johnson, 2014; Mitchell, Macrae, & Banaji, 2006), is posited to underlie idea generation, whereas the more goal-oriented executive control network (Fox et al., 2005; Zabelina & Andrews-Hanna, 2016) is thought to underlie idea evaluation (Beaty et al., 2015). While these two networks are typically anti-correlated (Fox et al., 2005, Zabelina & Andrews-Hanna, 2016), the flexible deployment of these dual neural and cognitive systems is linked to high creative ability (Zabelina & Robinson, 2010). Additionally, areas of the salience network may facilitate switching between default and executive processes (Sridharan, Levitan, & Menon, 2008; Uddin, 2015), and therefore may support this dynamic interplay between generative and evaluative processes during creative cognition (Beaty et al, 2018).

Returning to the behavioral model of improvisation (Pressing, 1988), the same networks thought to underlie generative and evaluative processes in domain-general creative cognition should also apply to the specific domain of musical improvisation. Previous work has already established that both DMN and ECN are involved in musical improvisation, with DMN showing increased activity during tasks in which participants were allowed to freely improvise (Limb & Braun, 2008; Liu et al., 2012), and ECN showing increased activity during tasks where improvisation was highly constrained in terms of available pitches or emotional character (Bengtsson, Csíkszentmihályi, & Ullén, 2014; Pinho et al., 2014). Previous work has also shown that task constraints can account for these differences in ECN and DMN activity (Pinho et al., 2015), with more constrained tasks showing more ECN activity and lower-constraint tasks showed more DMN activity. This is consistent with the dual-process theory, as higher task constraints could relate to a higher relevance of evaluative processes, and therefore increased activity of the ECN, whereas lower constraint tasks could rely more on generative processing by the DMN.

Given the alignment of behavioral and neural models of musical improvisation (Pressing, 1988; Loui, 2018), functional connectivity profiles associated with generalized creative behaviors (Beaty et al., 2015; 2018), and task fMRI studies of musical improvisation (Limb & Braun 2008; Liu et al., 2012; Bengtsson et al., 2014; Pinho et al., 2014), we expect to see differences in ECN and DMN in improvising musicians. In the present study, we ask whether improvisationally-trained musicians show different functional connectivity patterns in the absence of task, as has been observed in populations of individuals who are highly creative as measured by the Torrance test of creative thinking (TTCT) (Beaty et al., 2018; Takeuchi et al, 2012). Second, we sought to differentiate between the effects of improvisational and non-improvisational music training in resting state functional connectivity.

While there are improvisational components in many genres of music training, contemporary common-practice jazz music training places special emphasis on improvisation (e.g. Sawyer, 2014). In contrast, contemporary common-practice classical western music training places less emphasis on improvisation. Thus, we compared musicians who were primarily trained in jazz improvisation against musicians trained in the classical tradition, with the hypothesis that those with jazz improvisation training might show intrinsic brain networks that more resemble the real-time creative process of improvisation.

Up to this point, only one major task fMRI study has differentiated between improvisational and non-improvisational musical training (Pinho et al, 2014). However, multiple studies suggest that the same networks shown to be important for creative behaviors are also involved in music. Musical training has been associated with increased functional connectivity of the salience network (Luo et al., 2014) as well as increased executive functioning (Moreno et al., 2011) and heightened activity in the ECN (Sachs et al, 2017). Moreover, default mode activity has been associated with diverse musical behaviors, including the tracking of musical tonality (Janata et al., 2002), associating music with autobiographical memories (Janata, 2009), and the aesthetic response to music (Williams et al., 2018). Considering this, it is possible that differences in the functional connectivity profiles of improvisational musicians could be a result of general musical training rather than improvisational training in particular.

To disentangle the effects of musical improvisation and general musical training, here we compare resting state functional networks in improvisational musicians to those of musicians who do not improvise (hereafter classical musicians^1^), as well as minimally musically trained (MMT) controls. We use these groups as a means of differentiating between the impact of both presence and type (improvisational focused vs non-improvisational focused) of musical experience on resting state functional connectivity. While previous work has identified differences in resting state functional connectivity in musicians compared to non-musicians (Luo et al, 2012; Fauvel et al, 2014; Palomar-Garcia et al, 2017), in the present study we extend the sample to include two groups of musicians, and we examine results from the whole brain rather than focusing on auditory perceptual-motor networks as in previous studies. We focus first on comparing seed-based connectivity from the DMN and the ECN across the three groups. Then we relate these same networks to behavioral measures of musical improvisation and creativity tasks done outside the scanner. In regions that show between-group differences, we further localize the functional networks that gave rise to these differences and used those networks as seed regions of interest to compare their functional connectivity between groups, also relating the functional connectivity from those networks to creative behavior. Finally, we use tools from graph theory to compare network metrics across the groups, to assess the flexible deployment of neural systems across global and local networks that might underlie increased cognitive flexibility in more creative individuals (Zabelina & Robinson, 2010).

Our hypotheses are threefold: 1. We expect more functional connectivity from the DMN and ECN in both groups of musicians, compared to MMT controls, confirming general effects of musical training on functional connectivity of the brain as shown in previous studies (Luo et al, 2012; Fauvel et al, 2014; Palomar-Garcia et al, 2017). 2. We expect that seed regions in the DMN and ECN will be functionally connected to different out-of-network regions in Improvisational and Classical musicians, suggesting genre-specific effects of training. 3. We expect that Improvising musicians will have lower clustering and lower modularity than classical musicians, consistent with the previously observed pattern in creative individuals (Beaty et al, 2018), suggesting a more flexible organization of brain networks which facilitates the real-time constraints of musical improvisation.

## 2. Materials and Methods

### 2.1 Participants

Thirty-six participants were recruited from Wesleyan University and the Hartt School of Music and received monetary compensation or course credit for their participation. Participants were categorized into Improvisational, Classical, or Minimally Musically Trained (MMT) groups based on their reported musical experience. Due to difficulty recruiting a sufficiently large sample of female improvisational musicians to maintain gender-balanced groups, we limited our sample in all three groups to males.

The improvisational group (n = 12) was defined by the following criteria: 1) 5+ years of training in music that included improvisation, and 2) Current participation in improvisational musical activities for 1+ hours per week. The classical group (n = 12) was defined by the following criteria: 1) 5+ years of musical training, and 2) Current participation in non-improvisational musical activities for 1+ hour per week. The minimal musical training (MMT) group (n = 12) included participants who had less than 5 years of previous musical training.

The three groups were matched in age, general cognitive ability as measured by the Shipley abstraction test (Shipley, 1940), short term memory as measured by the digit span task (Baddeley, 2003), pitch discrimination threshold as measured by a staircase procedure (Loui et al., 2008), and age of onset of musical training (Table 1). Both the improvisational and classical groups had a longer duration of musical training than MMT, but there was no significant difference in duration or age of onset of musical training between the improvisational and classical groups. The jazz group had an average of 5.4 years of training (SD = 3.4 years) in musical improvisation, which was significantly higher than both the classical and MMT groups, who had 0.9 years (SD = 1.3 years) and 0 years of musical improvisation training respectively. Participants from the jazz group reported improvisational musical activities in one or more instruments including piano (n = 8), bass (2), guitar (3), mandolin (1), voice (1), drum (4), clarinet (1), saxophone (3), and vibraphone (1). Participants from the classical group reported non-improvisatory musical activities in piano (n = 2), bass (2), guitar (4), drum (3), clarinet (2), baritone (1), violin (1), saxophone (1), French horn (1), and pipa (1). All participants gave informed consent as approved by the Institutional Review Boards of Wesleyan University and Hartford Hospital.

**Table 1.** Performance on baseline behavioral tasks by group. For each task, the first number represents the mean value for the group, and the parenthetical number represents the standard deviation.

|  | Minimally Musically Trained (MMT) | Classical Musicians | Improvising Musicians | All Subjects |
| --- | --- | --- | --- | --- |
|  | (n=12 males) | (n=12 males) | (n=12 males) | (n=36 males) |
| Pitch Discrimination (Hz) | 7.8 (4.9) | 4.1 (2.4) | 4.1 (2.1) | 5.1 (3.5) |
| Digit Span (digits) | 7.5 (1.3) | 8.1 (1.9) | 7.9 (1.4) | 7.9 (1.5) |
| Shipley (raw score) | 17.7 (0.5) | 17.9 (1.5) | 17.3 (1.8) | 17.7 (1.4) |
| Age of onset of music training (years) | 9.8 (2.3) | 8.2 (3.4) | 8.4 (2.5) | 8.6 (2.8) |
| Duration of classical music training (years) | 0.9 (1.4) | 10.1 (3.7) | 8.4 (3.2) | 6.7 (4.9) |
| Duration of improvised music training (years) | 0 (0) | 0.9 (1.3) | 5.4 (3.4) | 2.3 (3.2) |

### 2.2 Procedures

After participants gave informed consent, they completed a questionnaire on their musical background, including questions about the age of onset of musical training, the duration of general musical training, and the duration of jazz and improvisation training (Table 1). A portion of our sample also performed two additional behavioral tasks that were used in exploratory follow-up brain-behavioral correlations. The two tasks are 1) a short version of the TTCT, (Torrance, 1968) (n = 18) and 2) the musical improvisation continuation task (ICT) measuring musical improvisation ability (Arkin et al., 2019) (n = 17). We include these behavioral tasks in exploratory follow-up analyses because previous work has shown that improvising musicians differ from classical musicians and MMT controls in their performance on the TTCT (Przysinda et al., 2017) and on the musical improvisation task (Arkin et al, 2019); thus we expect that the musical improvisation task and the TTCT provide useful measures of musical and domain-general creativity respectively.

#### 2.2.1 Improvisation Continuation Task

The Improvisation Continuation task was developed to assess musical improvisation in a laboratory setting, as a musical analog of the standard TTCT task (Arkin et al, 2019). Twelve different trials were presented from a computer. Each trial consisted of the presentation of a novel musical prompt (example audio files of the prompts are here: https://doi.org/10.6084/m9.figshare.6590489.v1). Participants were instructed to listen to the prompt (listening phase: two measures), then to play along with the prompt (continuation phase: eight measures), and then to improvise in the most creative way they could be based on the prompt (improvisation phase: eight measures). Visual cues were given on the screen throughout the listening phase (“Listen”), the continuation phase (“Play along”), and the improvisation phase (“Improvise”). A metronome was presented at 100 bpm to keep time throughout the entire trial. No additional instructions were given on how to improvise, nor were any guidelines given for how to play creatively. All participants completed this task on a Casio PX 150 MIDI keyboard. Participants who self-identified as playing other instruments additionally performed on their instrument of choice. Performances were recorded using a Zoom Q8 video camera.

#### 2.2.2 Torrance Test of Creative Thinking

Participants completed a short version of the TTCT (Torrance, 1968), in which they were given six open-ended verbal prompts (e.g. “List all the uses you can think of for a paper clip.”) and had three minutes to respond to each prompt. Participants were told that the task was a measure of general creativity and that they should try to give as many answers as they could.

#### 2.2.3 MRI Acquisition

High-resolution T1 and resting state images were acquired in a 3T Siemens Skyra MRI scanner at the Olin Neuropsychiatry Research Center at the Institute of Living. The anatomical images were acquired using a T1-weighted, 3D, magnetization-prepared, rapid-acquisition, gradient echo (MPRAGE) volume acquisition with a voxel resolution of 0.8 × 0.8 × 0.8 mm^3^ (TR = 2.4 s, TE = 2.09 ms, flip angle = 8º, FOV = 256 mm). Resting state MRI was acquired as 947 contiguous echo planar imaging (EPI) functional volumes (TR = 475 ms; TE = 30 ms; flip angle = 90, 48 slices; FOV = 240 mm; acquisition voxel size = 3 × 3 × 3 mm^3^). Participants kept their eyes open and fixated on a cross on the screen during resting state data acquisition. The resting state scan lasted 7.5 minutes.

### 2.3 Data Analysis

#### 2.3.1 Improvisation Continuation Task

Example audio files of participants’ production are available online at https://doi.org/10.6084/m9.figshare.6590489.v1. Audio was extracted from video recordings and rated for creativity by one professional jazz instructor (Rater 1) and one experienced jazz improvising musician (Rater 2), both of whom were blinded to the group status of each participant. Expert raters were asked to listen to each anonymized recording of each improvisation, and to rate the recording on a scale of 1–6 for creativity and imagination, with 1 being “Not creative and/or imaginative” to 6 being “Creative and/or imaginative.” Rater 1 completed 204 ratings (17 participants * 12 trials each), which included all participants in our sample that completed the task (9 improvising musicians, 6 classical musicians, and 2 MMT controls). Rater 2 completed ratings for 8 participants (6 trials each * 8 participants = 48 ratings), but stopped due to lack of interest. All available ratings across both raters were averaged for each participant (across either 12 or 18 trials), to obtain an averaged creativity rating for each participant.

#### 2.3.2. Torrance Test of Creative Thinking

Participants’ responses were coded for fluency and originality, as in previous studies in our lab (Przysinda et al, 2017). Fluency was calculated as the number of unique responses. Responses from 16 independent control participants (recruited online from Mechanical Turk) were used to create a baseline for originality. The participants were then scored for originality with unique responses receiving 3 points, responses that occurred once in the baseline receiving 2 points, and responses that occurred twice in the baseline receiving 1 point. Scoring of participants’ responses were done by raters who were blinded to the group status of each participant. Outlier trials were excluded using the Tukey method with a conservative multiplier of 2.2 (Tukey, 1977). This resulted in the removal of 2 single question scores in different participants, one from the Classical musician group and one from the Jazz group. Z-scores were calculated for each of the questions. In the current sample, fluency was highly correlated with originality (r = .92, p < .001), replicating previous studies (Kaufman, 2009). Since our previous work had shown that originality was a more effective measure than fluency at detecting differences between jazz and classical musicians (Przysinda et al, 2017), here we examined brain-behavior correlates with originality scores from the TTCT to avoid collinearity between behavioral predictor variables.

#### 2.3.3 MRI Preprocessing

Resting state and structural MRI preprocessing was carried out using the Statistical Parametric Mapping 12 (SPM12) software (http://www.fil.ion.ucl.ac.uk/spm/) with the CONN Toolbox (http://www.nitrc.org/projects/conn) (Whitfield-Gabrieli and Nieto-Castanon, 2012). In order, this consisted of functional realignment and unwarp, functional centering, functional slice time correction, functional outlier detection using the artifact detection tool (http://www.nitrc.org/projects/artifact_detect), functional direct segmentation and normalization to MNI template, structural centering, structural segmentation and normalization to MNI template, and functional smoothing to an 8mm gaussian kernel (Friston et al., 1995). Denoising steps for functional connectivity analysis included correction for confounding effects of white matter and cerebrospinal fluid (Behzadi et. al., 2007), and bandpass filtering to 0.008-0.09 Hz.

### 2.4 Seed-Based Connectivity Analyses

Since we were interested in whole-brain connectivity patterns of known whole-brain networks, we used a functional network atlas that related 14 cortical networks to known cognitive functions (Shirer et al., 2012). Mean time-course of each network was used as a single region-of-interest (ROI), and networks that resulted in clusters demonstrating a significant main effect of group were identified at the height threshold p < 0.001, uncorrected, and cluster size p < 0.05, p-FDR corrected level (Table 2). For each cluster that showed significant between-group differences, group-level beta values were extracted in order to determine the driving factors of these group differences. These networks were then used to extract group connectivity profiles at the height threshold p < 0.05, p-FWE corrected level. T-tests comparing each pair of groups were then performed for each network seed, and significant clusters were identified at the height threshold p < 0.001, uncorrected, and cluster size p < 0.05, p-FDR corrected level (Table 2). In order to relate connectivity differences directly to performance on creativity tasks, the creativity score on the ICT and originality score on the TTCT were used as second-level covariates. Clusters showing a significant correlation between task performance and network-seeded connectivity were identified at the height threshold p < 0.05, p-FWE corrected level. Anatomical locations for these clusters were identified using the Harvard-Oxford cortical and subcortical structural atlases (Makris et al., 2006; Frazier et al., 2005; Desikan et al., 2006; Goldstein et al., 2007).

**Table 2.**
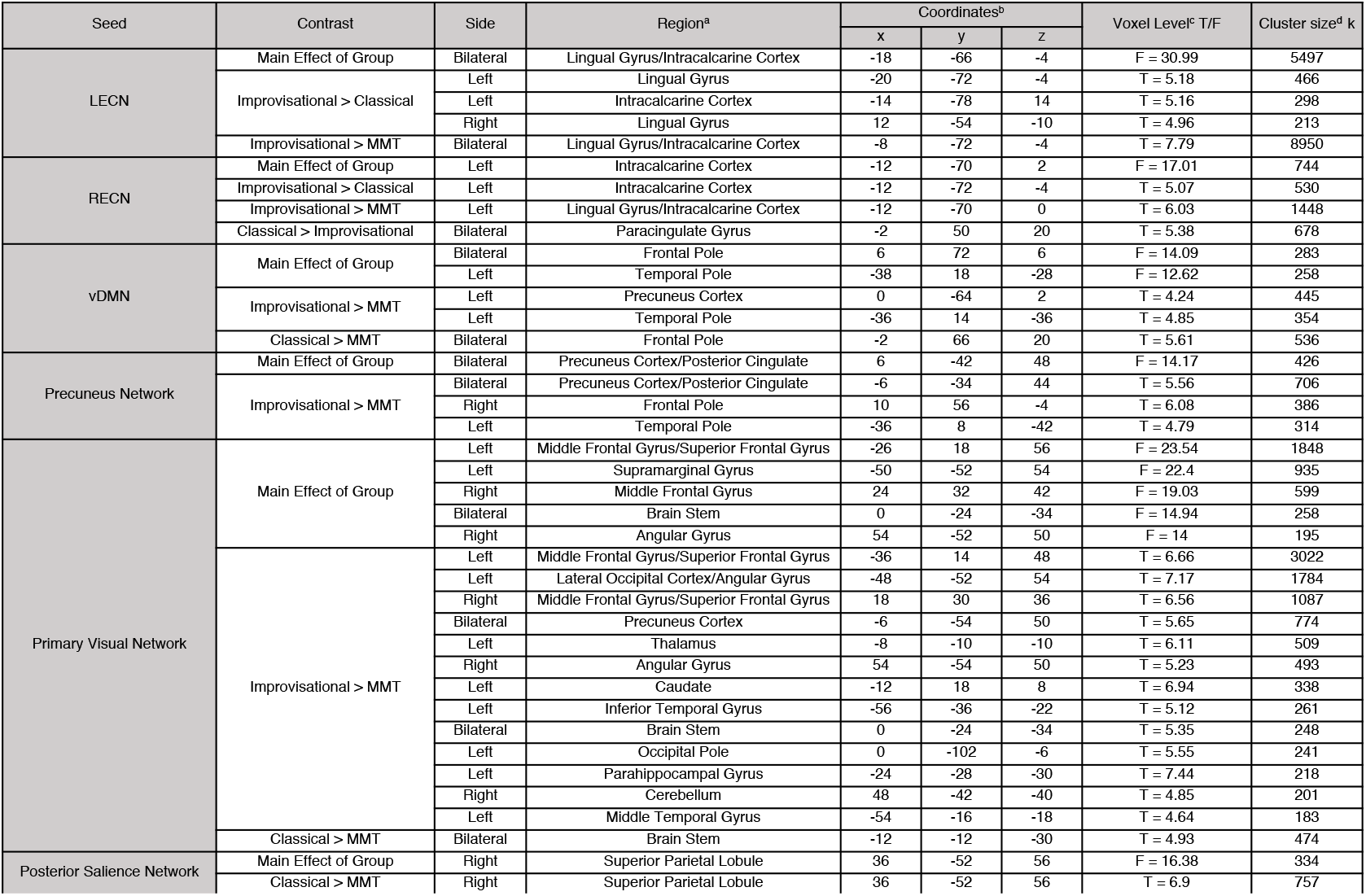
Brain regions showing connectivity differences related to musical experience. ^a^ Anatomical cluster locations indicated by Harvard-Oxford atlas. ^b^ Coordinates in millimeters in the Montreal Neurological Institute space. ^c^ Height threshold p < 0.001, uncorrected, one-sided contrast. ^d^ Cluster size p < 0.05, P-FWE corrected.

### 2.5 Graph Theory Analysis

Correlation matrices comparing all 90 regions of the 14-network Stanford atlas (Shirer et al., 2012) were extracted for each participant in each group. These matrices were then exported into MATLAB (mathworks.com) and analyzed using the Brain Connectivity Toolbox (Rubinov & Sporns, 2010). Each network statistic was computed at a range of correlation thresholds from r = 0.05 to r = 0.5. Individual participants’ measures of degrees, clustering coefficients, strengths, betweenness centrality, and local efficiency, were calculated for each region in each brain and then averaged across regions for each participant, whereas modularity was a single measure for the whole brain that was calculated for each participant. These group averages were then compared using one-way ANOVAs to determine group differences in each network measure while correcting for false-discovery rate of 0.05 for comparisons across 6 network measures (Benjamini-Hochberg, 1995).

## 3. Results

### 3.1 Seed-Based Connectivity Analysis

Significant main effects of group were observed in functional connectivity in both left and right ECN, as well as the ventral DMN, precuneus network, primary visual network, and posterior salience network (Table 2). Therefore, we limit our analysis of seed-based connectivity to these six networks.

#### 3.1.1 Left Executive Control Network

All groups showed highly significant left executive control network (LECN) functional connectivity to frontolateral and posterior parietal regions (Figure 1A), consistent with classically observed patterns of ECN connectivity (Fox et al., 2005). However, both musician groups showed larger areas of crossover into right hemispheric ECN connectivity than MMT controls (Figure 1A). Furthermore, the Improvisational group showed particularly high LECN connections to medial prefrontal cortex (mPFC) and posterior parietal cortex (PPC), suggesting heightened connectivity between ECN and DMN (Figure 1A). The Improvisational group also showed higher functional connectivity than both the Classical group and the MMT controls between LECN and bilateral lingual gyrus and intracalcarine cortex (Table 2; Figure 1B). In addition, LECN seeded connectivity correlated with both the creativity score on the ICT and the originality score on the TTCT within bilateral ECN areas, as well as mPFC and PCC from DMN and portions of the left temporal lobe and midbrain (Figure 1C).

**Figure 1.**
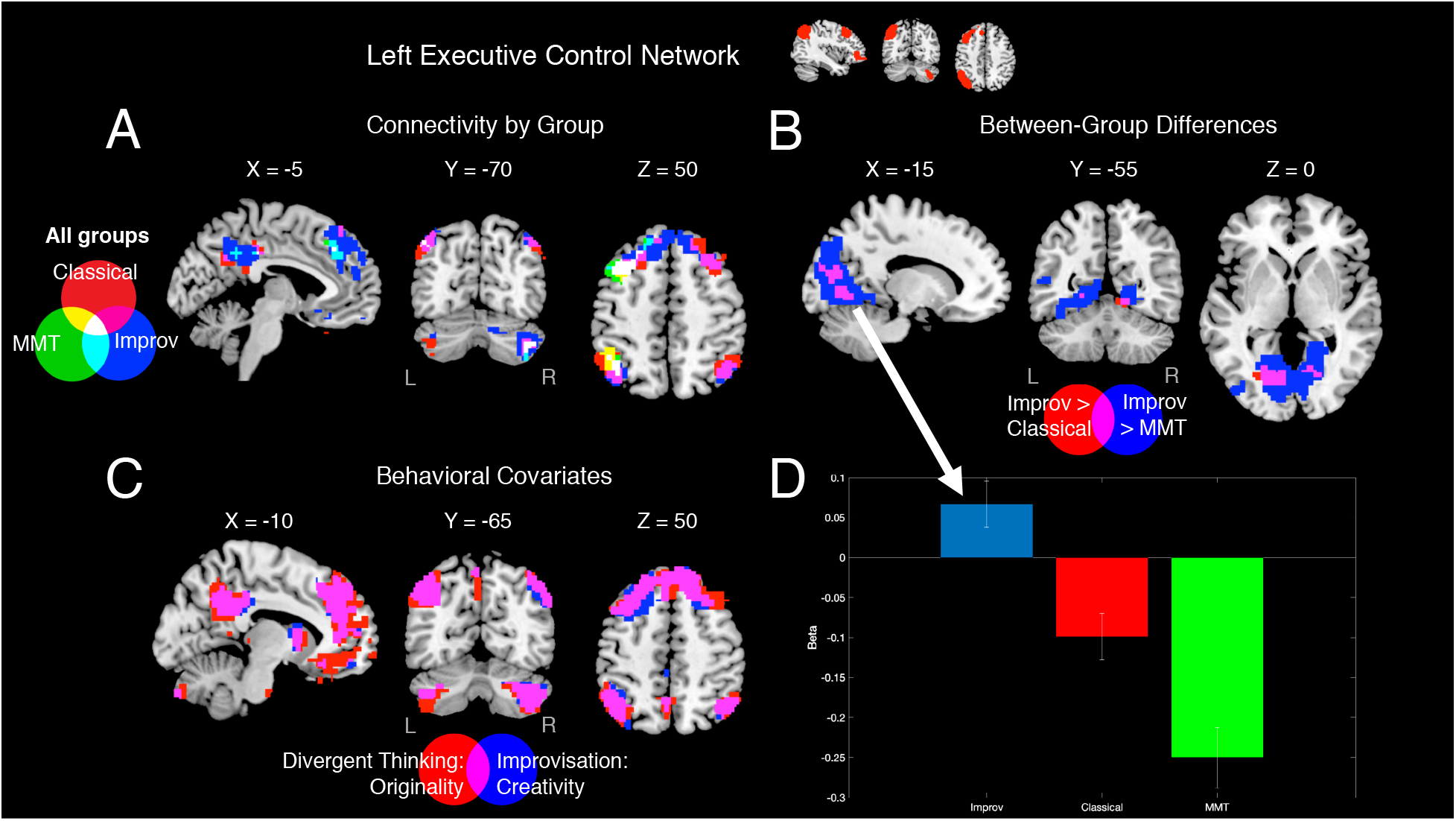
LECN Seed Based Connectivity Analysis. A: Connectivity profiles of improvisational group (blue), classical group (red), and MMT group (green) for LECN seed (height threshold p < 0.05, p-FWE corrected). B: Improvisational > Classical (red) and Improvisational > MMT (blue) group contrasts for LECN seed (height threshold p < 0.001, uncorrected, cluster size p < 0.05, p-FWE corrected). C: Regions demonstrating a significant correlation between LECN seeded connectivity and originality rating on the Torrance test of creative thinking (red) and creativity rating on the improvisation continuation task (blue) (height threshold p < 0.05, p-FWE corrected). D: Effect sizes for occipital cluster at (−18, −66, −4), showing a main effect of group on LECN seeded connectivity (bar = mean beta, error bar = between-subjects standard error).

#### 3.1.2 Right Executive Control Network

Connectivity profiles from the right executive control network (RECN) differed slightly from those observed in the LECN, especially in that differences that arose were higher for classical than for improvisational musicians. In this case, connectivity to PCC was present in both musician groups, and classical musicians also showed connectivity to bilateral temporal pole that was not seen in the other two groups (Figure 2A, coronal panel). as was the case with LECN, there was heightened connectivity between RECN and occipital areas in Improvisational musicians as compared to Classical musicians and MMT controls (Figure 2B/D, Table 2). The Classical group additionally showed higher connectivity than the Improvisational group from the RECN seed in portions of the paracingulate gyrus (Figure 2B, Table 2). Brain-behavior correlates indicated that the ICT and TTCT tasks related to functional connectivity between RECN and bilateral ECN areas, as well as DMN areas and portions of the midbrain and left cerebellum (Figure 2C).

**Figure 2.**
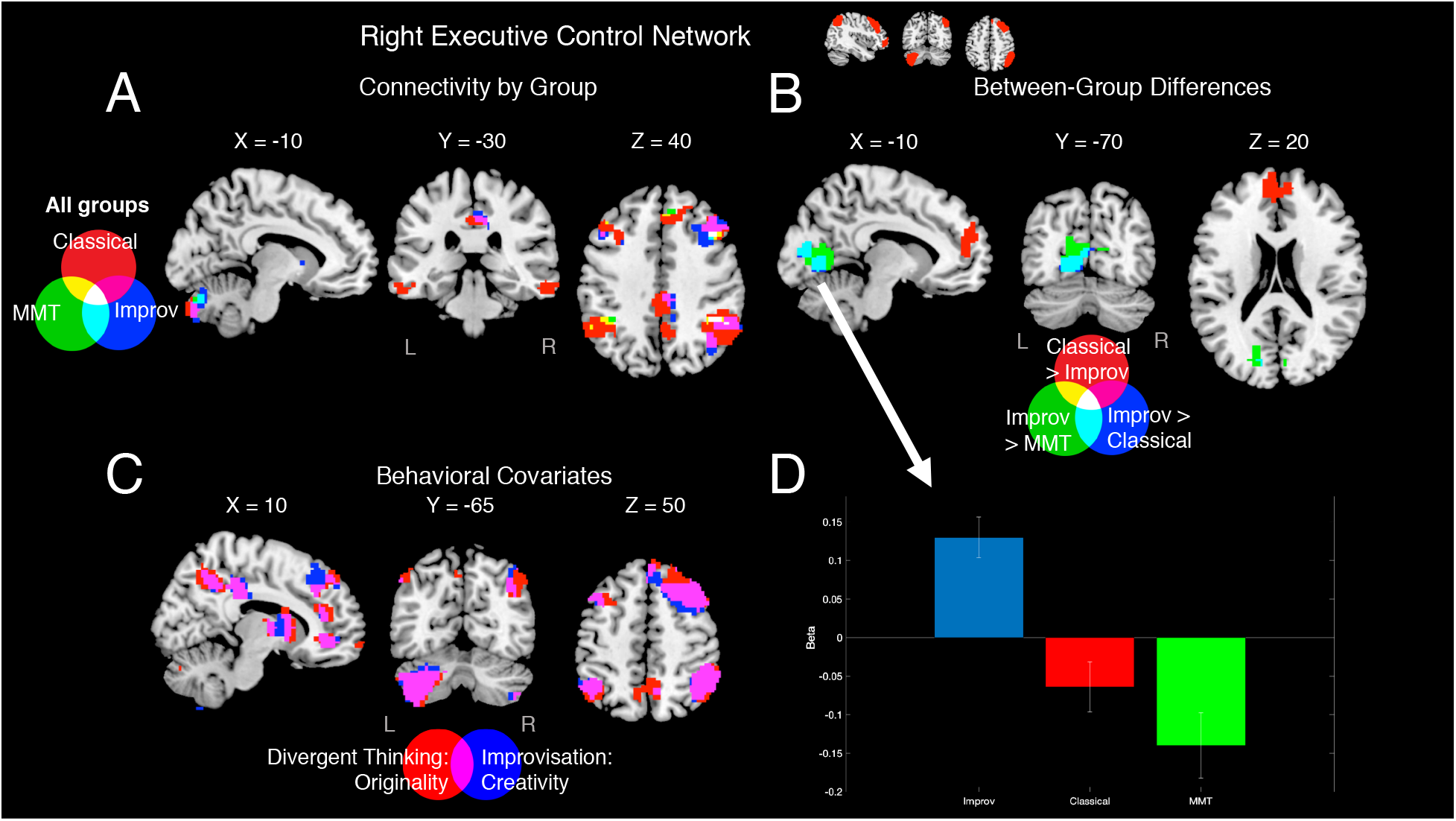
RECN Seed Based Connectivity Analysis. A: Connectivity profiles of improvisational group (blue), classical group (red), and MMT group (green) for RECN seed (height threshold p < 0.05, p-FWE corrected). B: Classical > Improvisational (red), Improvisational > Classical (blue) and Improvisational > MMT (green) group contrasts for LECN seed (height threshold p < 0.001, uncorrected, cluster size p < 0.05, p-FWE corrected). C: Regions that show a significant correlation between RECN seeded connectivity and originality rating on the Torrance Test of Creative Thinking (red) and creativity rating on the Improvisation Continuation Task (blue) (height threshold p < 0.05, p-FWE corrected). D: Effect sizes for occipital cluster (−12, −70, 2), showing a main effect of group on RECN seeded connectivity (bar = mean beta, error bar = standard error).

#### 3.1.3 Ventral Default Mode Network

While the ventral default mode network (vDMN) showed significant functional connectivity to PCC, consistent with the classic DMN, in all participants (Fox et al., 2005), the two musician groups showed larger clusters than MMT controls (Figure 3A). The Improvisational group showed higher functional connectivity than the MMT group in portions of precuneus cortex, intracalcarine cortex, and left temporal pole (Figure 3B/D, Table 2). In contrast, the Classical group showed higher connectivity than the MMT controls between vDMN and medial frontal pole (Figure 3B/D, Table 2). Performance on ICT and TTCT correlated with connectivity between vDMN and PCC, as well as some medial and lateral prefrontal regions and portions of bilateral cerebellum (Figure 3C).

**Figure 3.**
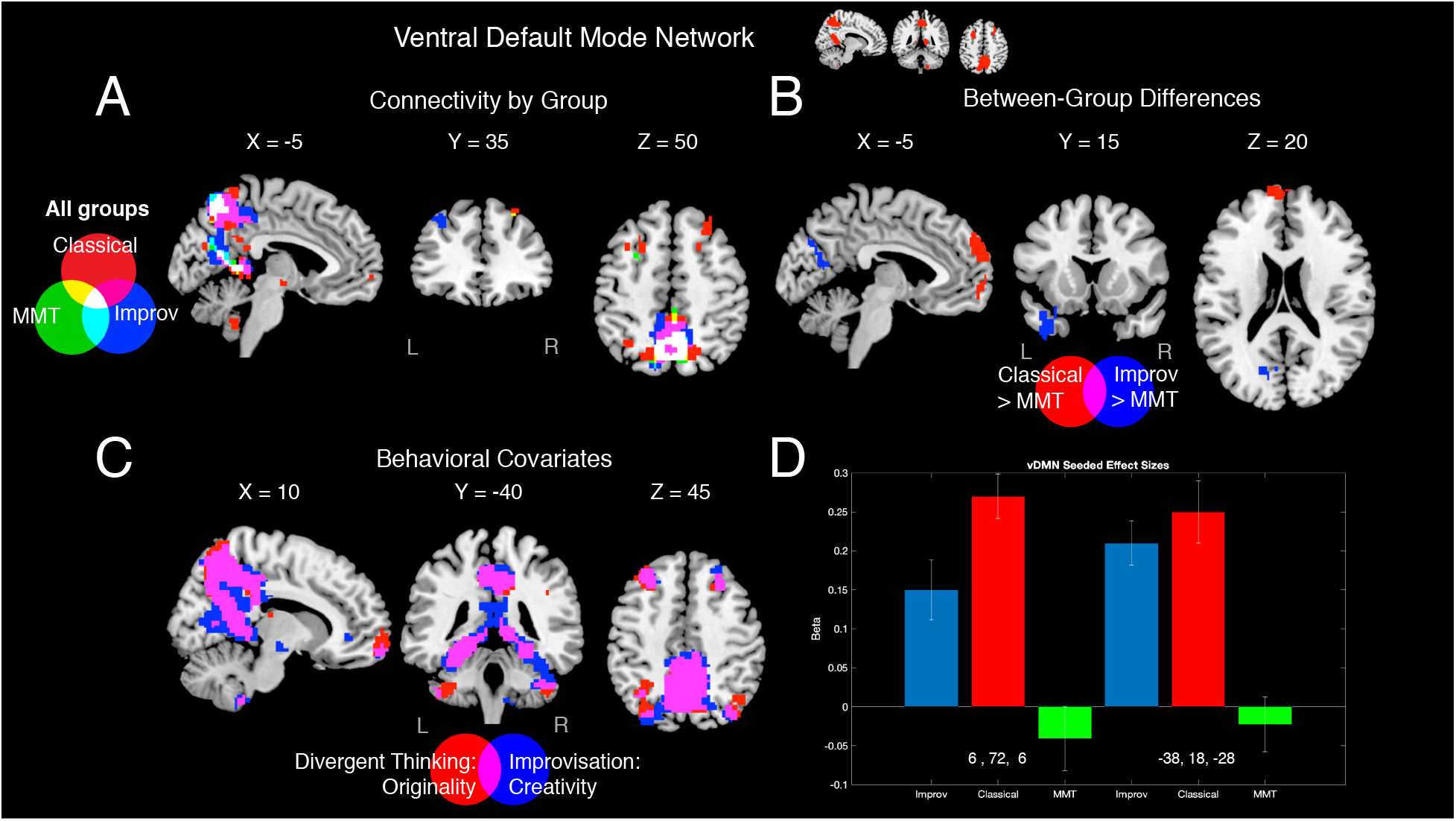
vDMN Seed Based Connectivity Analysis. A: Connectivity profiles of improvisational group (blue), classical group (red), and MMT group (green) for vDMN seed (height threshold p < 0.05, p-FWE corrected). B: Classical > MMT (red) and Improvisational > MMT (blue) group contrasts for vDMN seed (height threshold p < 0.001, uncorrected, cluster size p < 0.05, p-FWE corrected). C: Regions that show a significant correlation between vDMN seeded connectivity and originality rating on the Torrance test of creative thinking (red) and creativity rating on the improvisation continuation task (blue) (height threshold p < 0.05, p-FWE corrected). D: Effect sizes for vDMN clusters showing a main effect of group on vDMN seeded connectivity (bar = mean beta, error bar = standard error).

#### 3.1.4 Precuneus Network

Precuneal connectivity to PCC was especially high in the Improvisational group, and the improvisational group also showed connectivity to medial frontal regions not seen in the other two groups (Figure 4A). This difference between groups was significant when comparing the Improvisational group to MMT controls, with significant clusters found in the PCC and medial frontal pole (Figure 4B/D, Table 2). Furthermore, the Improvisational group showed higher Precuneus network seeded functional connectivity than the MMT group in the portions of the left temporal pole (Figure 4B, Table 2). A large cluster including PCC and a significant portion of the occipital lobe showed a correlation between precuneus network connectivity and performance on the ICT and TTCT, in addition to the medial prefrontal cortex from DMN and portions of lateral prefrontal cortex from ECN (Figure 4C).

**Figure 4.**
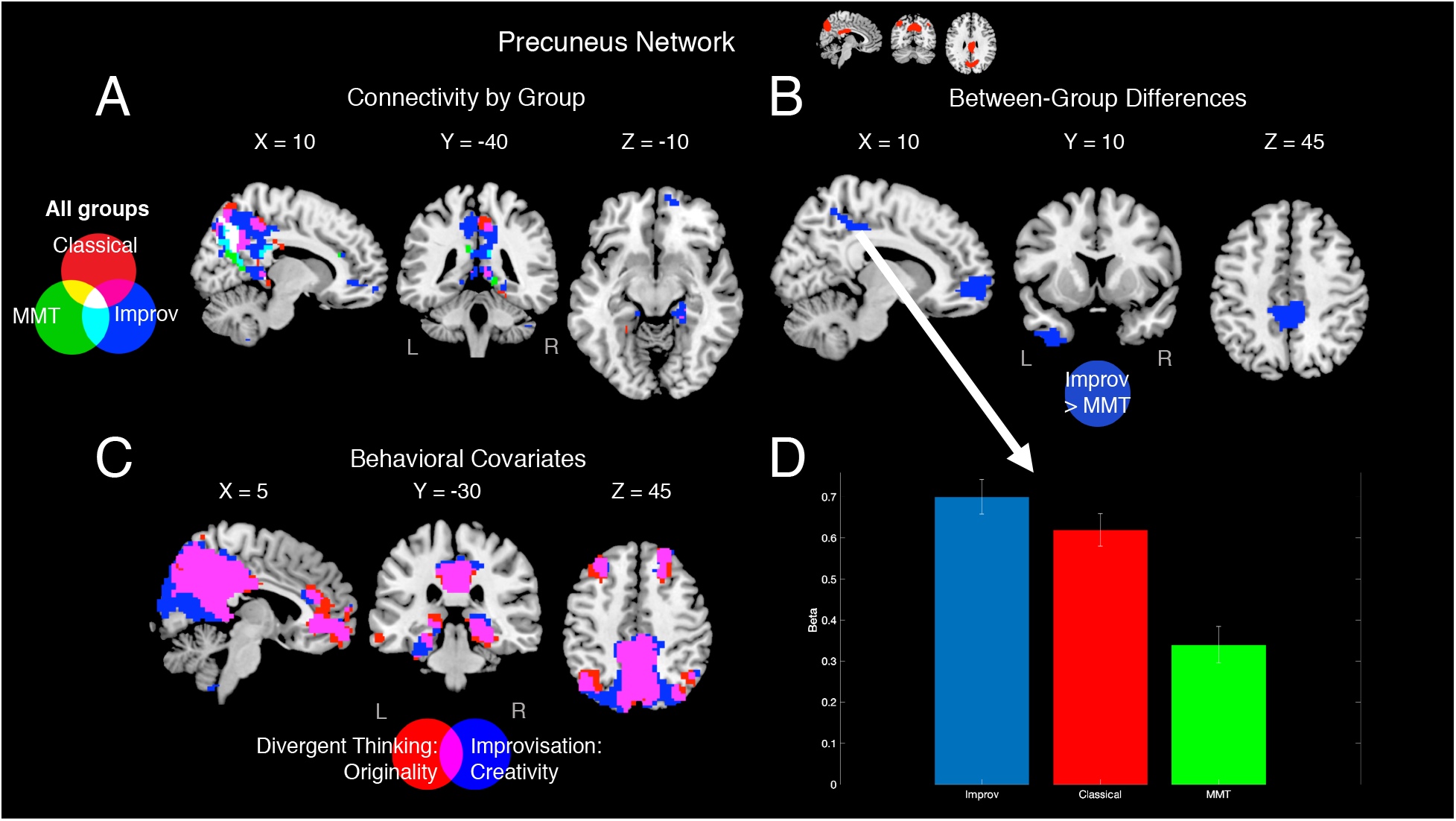
Precuneus Network Seed Based Connectivity Analysis. A: Connectivity profiles of improvisational group (blue), classical group (red), and MMT group (green) for precuneus network seed (height threshold p < 0.05, p-FWE corrected). B: Improvisational > MMT group contrast for precuneus network seed (height threshold p < 0.001, uncorrected, cluster size p < 0.05, p-FWE corrected). C: Regions that show a significant correlation between precuneus network seeded connectivity and originality rating on the Torrance test of creative thinking (red) and creativity rating on the improvisation continuation task (blue) (height threshold p < 0.05, p-FWE corrected). D: Effect sizes for precuneal cluster (6, −42, 48), demonstrating a main effect of group on precuneus network seeded connectivity (bar = mean beta, error bar = standard error).

#### 3.1.5 Primary Visual Network

Group connectivity profiles for primary visual network were largely confined to the occipital lobe, except in the improvisational group, which included significant connections to frontal and midbrain regions (Figure 5A). The Improvisational group showed higher functional connectivity than the MMT group in occipital pole, lateral occipital cortex, bilateral angular gyrus, precuneus cortex, left middle and inferior temporal gyrus, left parahippocampal gyrus, bilateral superior and middle frontal gyrus, bilateral thalamus, left caudate nucleus, and portions of the right cerebellum and the brainstem (Figure 5B/D, Table 2). Classical musicians also showed higher connectivity than MMT controls between primary visual network and the brainstem (Figure 5B/D, Table 2). Primary visual network connectivity was significantly correlated with performance on the ICT and TTCT for a large portion of the medial occipital lobe and portions of the medial cerebellum, as well as some small portions of the frontal and parietal lobes similar to what was seen in the Improvisational group (Figure 5C).

**Figure 5.**
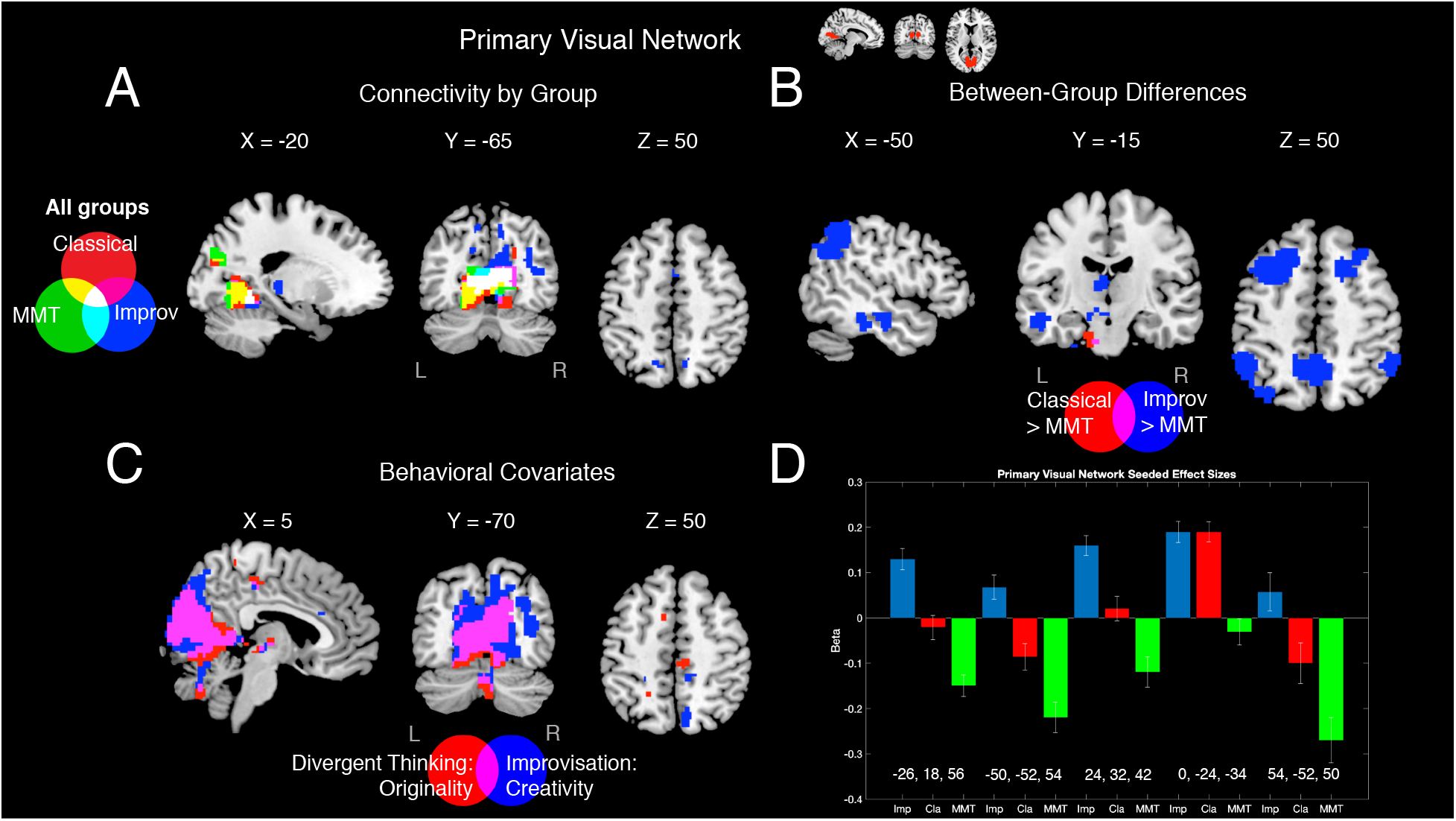
Primary Visual Network Seed Based Connectivity Analysis. A: Connectivity profiles of improvisational group (blue), classical group (red), and MMT group (green) for primary visual network seed (height threshold p < 0.05, p-FWE corrected). B: Classical > MMT (red) and Improvisational > MMT (blue) group contrasts for primary visual network seed (height threshold p < 0.001, uncorrected, cluster size p < 0.05, p-FWE corrected). C: Regions that show a significant correlation between primary visual network seeded connectivity and originality rating on the Torrance test of creative thinking (red) and creativity rating on the improvisation continuation task (blue) (height threshold p < 0.05, p-FWE corrected). D: Effect sizes for clusters demonstrating a main effect of group on primary visual network seeded connectivity (bar = mean beta, error bar = standard error).

#### 3.1.6 Posterior Salience Network

Group connectivity profiles for posterior salience network were notably different among the two musician groups. Improvisational musicians showed larger clusters of connectivity to bilateral insular cortex and PCC than the other two groups, and classical musicians showing a cluster of connectivity to right superior parietal lobule not seen in the other groups (Figure 6A). However, group contrasts indicated that only this last cluster was significant, with the effect primarily driven by contrasts between Classical group and MMT controls (Figure 6B/D, Table 2). Lateral parietal and frontal regions, ventral cerebellum, and portions of the midbrain all showed significant correlation between posterior salience network and performance on the ICT and TTCT (Figure 6C).

**Figure 6.**
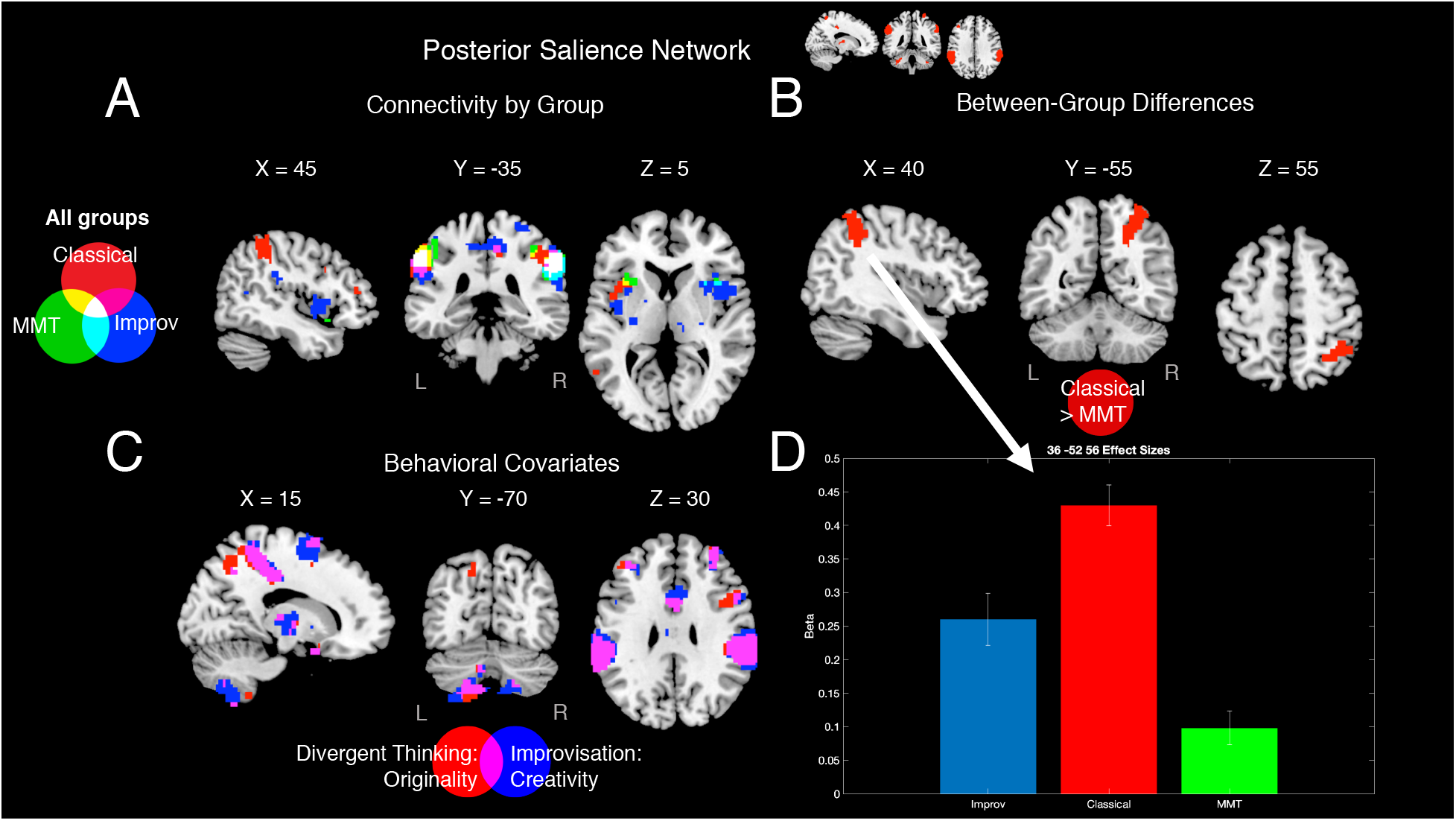
Posterior Salience Network Seed Based Connectivity Analysis. A: Connectivity profiles of improvisational group (blue), classical group (red), and MMT group (green) for posterior salience network seed (height threshold p < 0.05, p-FWE corrected). B: Classical > MMT group contrast for posterior salience network seed (height threshold p < 0.001, uncorrected, cluster size p < 0.05, p-FWE corrected). C: Regions that show a significant correlation between posterior salience network seeded connectivity and originality rating on the Torrance test of creative thinking (red) and creativity rating on the improvisation continuation task (blue) (height threshold p < 0.05, p-FWE corrected). D: Effect sizes for cluster demonstrating a main effect of group on posterior salience network seeded connectivity (bar = mean beta, error bar = standard error).

### 3.2 Graph Theory Analysis

Network measures of *degree* (the number of nodes significantly correlated to a given node), *strength* (the sum of the correlation coefficients for a given node), and *modularity* (the degree to which a network segregates into independent modules) did not differ between groups. However, several network measures showed main effects of groups; in each case, the measure was highest in the Classical group, followed by the Improvisational group and MMT controls, respectively. There was a significant main effect of group at a correlation threshold of r = 0.2 on *betweenness centrality*, (F(2, 267) = 5.95, p = 0.006, Benjamini-Hochberg corrected), which is the number of shortest paths from one node to another that contains a given node (Figure 7A), with classical musicians showing the highest betweenness centrality, followed by improvisational musicians and then by MMT. Similarly, *clustering coefficient*, the fraction of nodes correlated with a given node that are also correlated with one another, also showed a main effect of group, and was again highest in classical musicians (F(2, 267) = 13.02, p = 0.000004, Benjamini-Hochberg corrected, Figure 7B). *Local efficiency*, the average connectedness in the neighborhood of a given node, also showed a main effect of group, with the same relative pattern of classical musicians being highest (F(2, 267) = 8.41, p = 0.0003, Benjamini-Hochberg corrected, Figure 7C).

**Figure 7.**
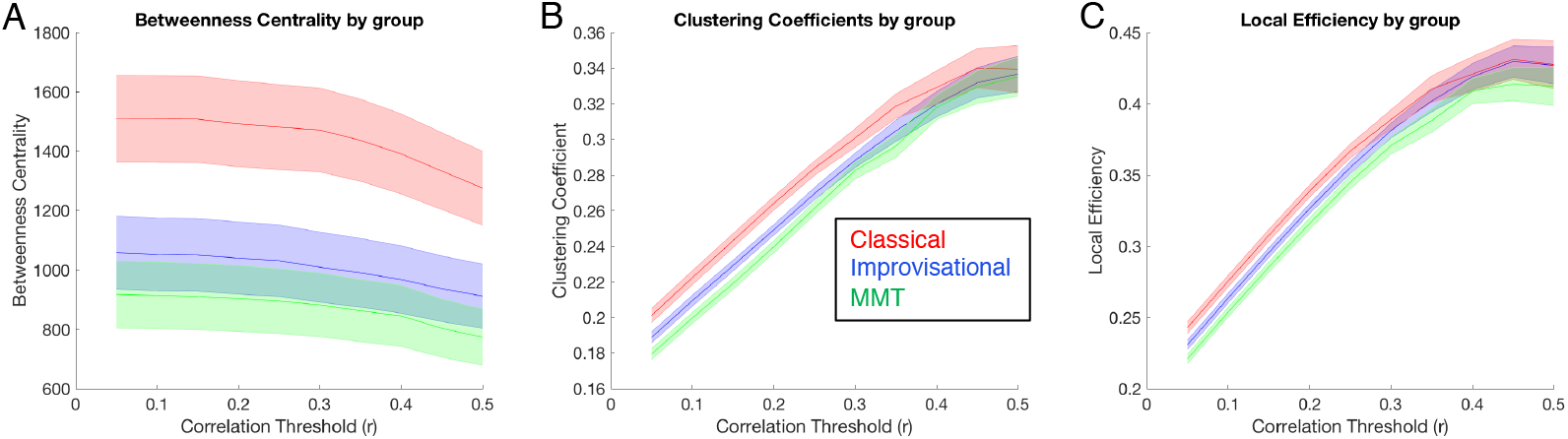
Group Differences in Small World Brain Connectivity. A: Betweenness centrality for classical musicians (red), improvisational musicians (blue), and MMT controls (green) across a range of correlation thresholds (solid line = mean of all participants/ROI’s for each group, error bar = standard error for all 90 Stanford ROI’s averaged across participants for each group). B: Clustering coefficients for classical musicians (red), improvisational musicians (blue), and MMT controls (green) across a range of correlation thresholds (solid line = mean of all participants/ROI’s for each group, error bar = standard error for all 90 Stanford ROI’s averaged across participants for each group). C: Local efficiency for classical musicians (red), improvisational musicians (blue), and MMT controls (green) across a range of correlation thresholds (solid line = mean of all participants/ROI’s for each group, error bar = standard error for all 90 Stanford ROI’s averaged across participants for each group).

## 4. Discussion

Creativity has been related to functional connectivity in the DMN and ECN, but how the interplay between these and other networks in the brain might give rise to musical creativity is unknown. Here we operationalize real-time musical creativity as training in musical improvisation, and we identify intrinsic brain networks that relate to improvised and non-improvised musical training. Results show generally increased functional connectivity in both musically-trained groups (both improvising and non-improvising) from both the DMN and the ECN as measured by seed-based connectivity analyses.

### 4.1 Differences in Resting State Connectivity Related to General Music Training

Seed based group connectivity profiles for ECN and DMN regions showed a uniform pattern of higher functional connectivity in both musician groups as compared to MMT controls. We also saw some musical training-related increases in salience network functional connectivity, as has been previously reported (Luo et al., 2014). Where connectivity differences related to general musical training did exist, they tended to be in one of two forms: within-network connectivity and between-network connectivity.

The first set of findings were differences in intrinsic within-network connectivity. That is, group profiles of DMN seeds tended to show more connectivity to other DMN regions, and ECN seeds tended to show more connectivity to other ECN regions within the musician groups. In particular, the two musician groups showed higher RECN seeded connectivity to LECN and higher LECN seeded connectivity to RECN. This adds support for previous work showing effects of musical training on higher executive function (Sachs et al, 2017). Additionally, Classical musicians showed significantly higher connectivity than the MMT group between vDMN and the mPFC, which is a known hub of the DMN. This result suggests that classical musical training is linked to more within-network connectivity, especially within the DMN.

The second set of findings pertain to between-network connectivity. While most of the group contrasts with the primary visual network seed were higher in the Improvisational group than in MMT controls, there was one cluster of brainstem connectivity that was higher in both musician groups relative to MMT controls. Furthermore, superior temporal lobe structures in musicians showed different connectivity patterns from MMT controls: Classical musicians had significant connectivity from RECN to bilateral temporal pole which was not seen in the other two groups, whereas Improvisational musicians had higher connectivity than MMT controls between vDMN seed and left temporal pole, precuneus network seed and left temporal pole, and primary visual network seed and left middle and inferior temporal gyri. The latter is consistent with magnetoencephalography work showing that musicians have more densely connected cortical network that enables audiovisual integration (Paraskevopoulos et. al., 2015). Together, our results suggest that while classical musicians may recruit auditory cortex more for executive processes, improvisational musicians may recruit auditory cortex for more default and primary sensory functioning.

### 4.2 Differences in Resting State Connectivity Related to Improvisation-Specific Music Training

While some of the seed-based connectivity results reflected an association between DMN and ECN connectivity and general music training, there were also other striking differences between Classical and Improvisational group in resting state connectivity. Firstly, improvisational and classical musicians differ in connectivity from the primary visual network. The primary visual network seed showed many differences between the Improvisational group and MMT controls, and bilateral ECN seeds each showed higher connections to visual areas including intracalcarine cortex and lingual gyrus in Improvisational musicians relative to the other two groups. Further examining the effect sizes of each group confirmed that Improvisational musicians had high functional connectivity from primary visual areas to regions where MMT controls had negative beta values, or anticorrelations in these regions as they classically belong to different networks (Figure 5D). The finding of higher occipital or visual network connectivity in improvising musicians is consistent with an emerging body of work in EEG, DTI, and VBM associating occipital lobe connectivity with domain-general creativity (Petsche, 1996; Takeuchi et al., 2010; Zamm et al, 2013; Fink et al., 2014). Takeuchi et al. (2010) found that white matter in the lingual gyrus were correlated with performance on a creative task. Zamm et al (2013) found that people with music-color synesthesia, who were more likely to engage in creative industries, had higher white matter connectivity in the inferior frontal-occipital fasciculus, specifically in the fusiform gyrus. Fink et al (2014) found that grey matter density within the occipital lobe was correlated with verbal creativity. While visual areas are not a part of the high creative network (cf. Beaty et al, 2018), this agreement across a breadth of methodological approaches leads us to the possibility that highly connected visual cortices may underlie domain-general creativity, possibly by facilitating creative imagery.

However, not all differences between Improvisational and Classical Group connectivity showed higher connectivity in the former relative to the latter. In the vDMN seed, Improvisational musicians showed higher connectivity to Precuneus cortex, whereas Classical musicians showed higher connectivity to frontal pole. This suggests a more dominant prefrontal DMN component in classical musicians. Furthermore, Classical musicians showed higher connectivity than Improvisational musicians between RECN and paracingulate gyrus, suggesting that frontal executive connectivity is more relevant to classical than to improvisational musicianship. Finally, connectivity differences between posterior salience network and superior parietal lobule (SPL) were driven by differences between classical musicians and MMT controls. SPL has been associated with selective attention and memory retrieval during music listening (Satoh et al., 2001; Klostermann, Loui, & Shimamura, 2009), and it follows that classical musicians would be more capable of finely focusing their attention and activating retrieval mechanisms than improvisational musicians who must remain attentive to all aspects of a performance.

Moving now to graph theoretical analyses, Classical musicians showed highest betweenness centrality, clustering coefficients, and local efficiency amongst the three groups. This suggests that there are fewer, more tightly-knit communities in classical musicians, contrasting with more large-scale, globally-connected networks in improvisational musicians. This is further evidenced by the fact that where between-group differences in seed-based connectivity were observed, the Improvisational group consistently showed positive beta values, often contrasting with negative values in one or both of the other groups. This lack of tightly knit communities in improvisational musicians, coupled with more interactions between ECN and DMN, corresponds well with expected connectivity patterns of improvisational musicians: As improvisation requires constant retrieval of previously-stored musical knowledge, through which musical goals and semantic referents (e.g. musical themes) are filtered, in order to generate auditory-motor sequences that are novel and yet closely integrated with past knowledge (Pressing 1998, see also Loui, 2018).

### 4.3 Behavioral Correlates of Functional Connectivity

Across all six seed networks, behavioral performance on the TTCT and ICT correlated strongly with connectivity of the network in question. Additionally, there was a strong correlation between task performance and connectivity between ECN seeds and DMN regions, between vDMN seed and both visual and ECN regions, and between Precuneus network seed and DMN, ECN, and visual network regions. Notably, these clusters of connectivity tended to be very similar between the two different tasks, despite the fact that the TTCT and ICT are overtly very different, with the TTCT requiring written verbal responses while the ICT required music performance. This suggests a strong connection between general creativity, musical improvisation, and functional connectivity across the cortical networks most critical to creative cognition.

### 4.4 Limitations and Future Directions

While these findings are fairly consistent with our hypotheses and the existing body of literature, there are certain limitations to our sample and experimental procedures that should be addressed. Previous work has pointed out that despite no differences in standardized tests of creativity, women are less likely to enter “highly creative” fields such as musical improvisation, likely due to sociocultural reasons as jazz is still a very male-dominated field (Baer & Kaufman, 2008). Within our sampling pool, we were unable to recruit a sufficient sample of female improvisational musicians. While previous EEG studies comparing classical and jazz musicians have used unequal gender distributions between the groups (Bianco et al, 2018), previous MRI work has shown gender effects in functional connectivity (Schmithorst & Holland, 2006; Biswal et al., 2010; Zuo et al., 2010) and in structural connectivity as it relates to creativity (Ryman et al., 2014). These findings led us to avoid a gender confound within our sample by excluding female participants from Classical and MMT groups, which in turn affected our sample size for behavioral data.

Future plans include recruiting larger samples of both male and female participants, and, importantly, better determining causality in the association between brain connectivity and different forms of musical training through a longitudinal study on participants over the course of their musical training. Finally, we would also be interested in exploring the impact of other modes of musical creativity on brain connectivity. In particular, musical composition has the same end result of a novel and appropriate musical product, but achieves this on a much slower time-scale than musical improvisation. This may result in a greater involvement of evaluative processes, as well as less dynamic interplay between generative and evaluative processes. Therefore, comparisons of improvisational and compositional musicianship may prove useful in better characterizing the neural networks underlying musical creativity.

### 4.5 Conclusions

Intrinsic functional networks of improvisational musicians show more global resting state connectivity across executive control, default mode, and primary visual networks. This is consistent with previous literature suggesting that these networks are critical for the dynamic interplay of generative and evaluative processes (Beaty et al., 2015; Beaty et al., 2018; Petsche, 1996) and imagery associated with creative ideation (Takeuchi et al., 2010; Fink et al., 2014). In contrast, classical musical training is associated with increased within-network resting state connectivity of the DMN and the ECN. Results distinguish for the first time between effects of improvisational and non-improvisational musical training on functional connectivity, even in the absence of task.

## Data and Code Availability Statement

Group functional connectivity maps are publicly available on Neurovault: https://neurovault.org/collections/XVPXULLL/

Additional data, and code that support the findings of this study, are available from the corresponding author upon reasonable request.

## Footnotes

1 Non-improvising musicians in this study are mostly trained in the classical tradition. This is not to say that classical musicians do not improvise, as improvisation has historically been an important part of classical music training, e.g. in figured bass realization. However, the modern traditional classical training institutions, from which we recruit our participants, do not routinely provide training in improvisation; thus we adopt the “classical” label for our non-improvising musicians for the purposes of this study.

---

**Supplemental Figure S1.**
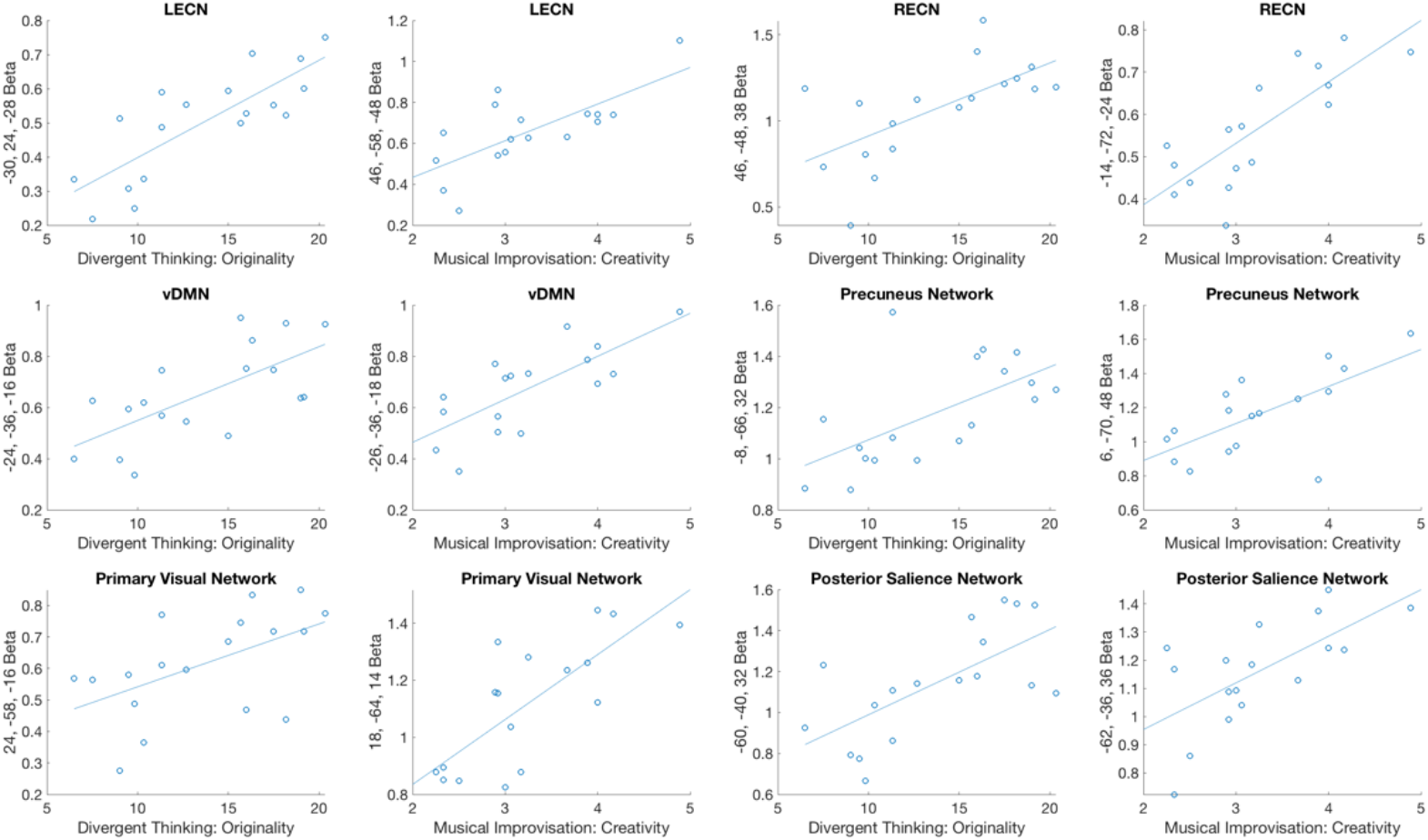
Correlations Between Creative Behavior and Network Seeded Connectivity. Beta values for global maximum of network seeded effect of performance on divergent thinking and musical improvisation tasks (Y axes) plotted against performance on these tasks (X axes) for each participant.

